# Biosynthesis of Kratom Opioids

**DOI:** 10.1101/2022.12.25.521902

**Authors:** Kyunhee Kim, Mohammadamin Shahsavarani, Jorge Jonathan Oswaldo Garza-García, Jack Edward Carlisle, Jun Guo, Vincenzo De Luca, Yang Qu

## Abstract

*Mitragyna speciosa* (kratom) derived monoterpenoid indole alkaloids (MIAs) such as mitragynine and 7-hydroxymitragynine are a new class of opioids with a corynanthe MIA pharmacophore that is responsible for their significantly reduced side effects and superior safety profiles. While botanical kratom has been historically used for stimulation and pain management in Southeast Asia, the biosynthesis of kratom MIAs is not known. In this study, we identified and characterized 9 reductases bearing various degrees of demethyldihydrocoryanthine/demethylcorynantheidine synthase activity and a new SABATH type methyltransferase that catalyzes highly unusual non-aromatic enol methylation from kratom and several other species, which are required in kratom opioids biosynthesis. With unnatural substrate 4-hydroxytryptamine, we further showed the biosynthesis of mitragynine and its epimer speciogynine using these characterized enzymes. The promiscuity of kratom opioid biosynthetic enzymes suggests that derivatives and analogs of kratom opioids may be manufactured in heterologous systems with appropriate enzymes and substrates.

## Introduction

Mitragynine from the plant *Mitragyna speciosa* (Kratom) and its derivatives and semi-synthetic analogs including 7-hydroxymitragynine and mitragynine pseudoindoxyl are new opioids with superior side effect profiles ^1,2^. Comparing to the widely used opioids such as morphine, these monoterpenoid indole alkaloid (MIA) based opioids have distinct opioid receptor agonism and signal transduction pathway, which does not recruit β-arrestin-2 that is linked with respiration depression ^1–5^. In animal models, kratom opioids did not promote self-administration ^6^, suggesting that it will less likely lead to addiction, one of the fundamental causes for the devastating opioid overdose epidemic around the world. Recently, the structural activity relationship (SAR) studies have shown that the scaffold of kratom opioids is amenable to modifications and substitutions, which could further improve the drug safety^1,5^. While kratom have been historically used as an ethnobotanical remedy for hundreds of years in Southeast Asia and the total synthesis has been reported ^2^, the biosynthesis of kratom opioids from the central precursor strictosidine to over 3,000 MIAs is not known. In this study, we report the discoveries and characterizations of nine reductases and the C17-enol methyltransferase that are responsible for the biosynthesis of mitragynine and related corynanthe alkaloids in kratom and other plant species. With the unnatural substrate 4-hydroxytryptamine, we further show the biosynthesis of mitragynine and its 20-epimer speciogynine using the characterized enzymes.

## Results

Previously our group and others have characterized the critical reductase geissoschizine synthase (CrGS) from the plant *Cathranthus roseus* (Madagascar’s periwinkle) ^7–9^. CrGS reduces the strictosidine aglycones to the corynanthe type MIAs 19*E*- and 19*Z*-geissoschizine (*m/z* 353, 19,20-dehydro, **3, 4**), the former of which is intermediate to major MIA skeletons including strychnos, sarpagan, iboga, and aspidosperma (Fig. 1). In addition to geissoschizine, the heteroyohimbine MIAs such as ajmalicine (**5**), tetrahydroalstonine (THA, **6**), and mayumbine also derive from reducing strictosidine aglycones by homologous reductases such as heteroyohimbine synthase (CrHYS) and tetrahydroalstonine synthase (CrTHAS1-4) ^10^. From the plant *Cinchona puberscens*, Trenti et al. characterized the reduction of strictosidine aglycones to demethyldihydrocorynantheine (*m/z* 355, corynanthe type, 20*S*, **1**) by another homologous reductase demethyldihydrocorynantheine synthase (CpDCS) ^11^, which is intermediate to the antimalarial drug quinine (Fig. 1). The biosynthesis of mitragynine (**18**) instead involves the formation of demethylcorynantheidine (**2**), the 20*R*-epimer of demethyldihydrocorynantheine (**1**). The remaining steps include *O*-methylation on the 17-enol, hydroxylation at C9, and finally *O*-methylation at 9-hydroxyl (Fig. 1). In addition to mitragynine and its intermediates, kratom and other related Rubiaceae species also make a number of stereo- and geometric isomers differing at C3, C15, and C18-19-20 (Fig. 1) ^4,12^. These diverse structures, such as specioginine (20*S*, **17**), speciociliatine (3*R*, **19**), and paynantheine (20*S*, 18,19-dehydro, **20**), suggest that a number of reductases are likely involved in their making.

**Figure 1.**
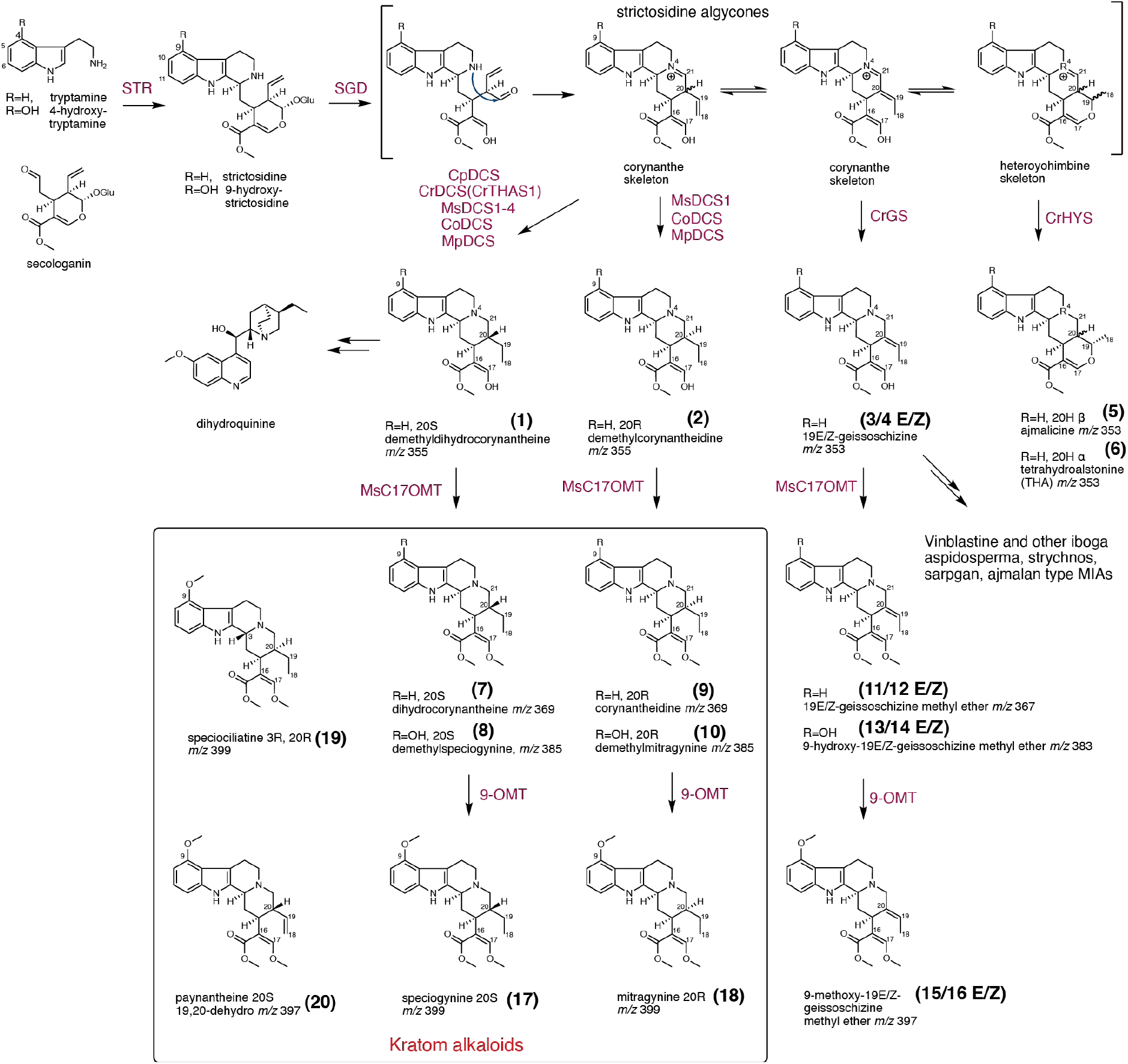
The biosynthetic pathway for kratom alkaloids and other related monoterpenoid indole alkaloids.

With the sequences of CrGS, CrTHAS1, and CpDCS, we identified and cloned four homologous reductases from the *M. speciosa* leaf transcriptome, which were named as demethyldihydrocorynantheine/demethylcorynantheidine synthases 1-4 (MsDCS1-4). MsDCS1 has been previously cloned by Trenti et al. (Genbank MW456555), however the gene product was not characterized. The four MsDCS shared 80-95% identity among each other, and were clustered together from other characterized of short/medium chain alcohol dehydrogenases in a phylogenetic analysis (Fig. 2a).

**Figure 2.**
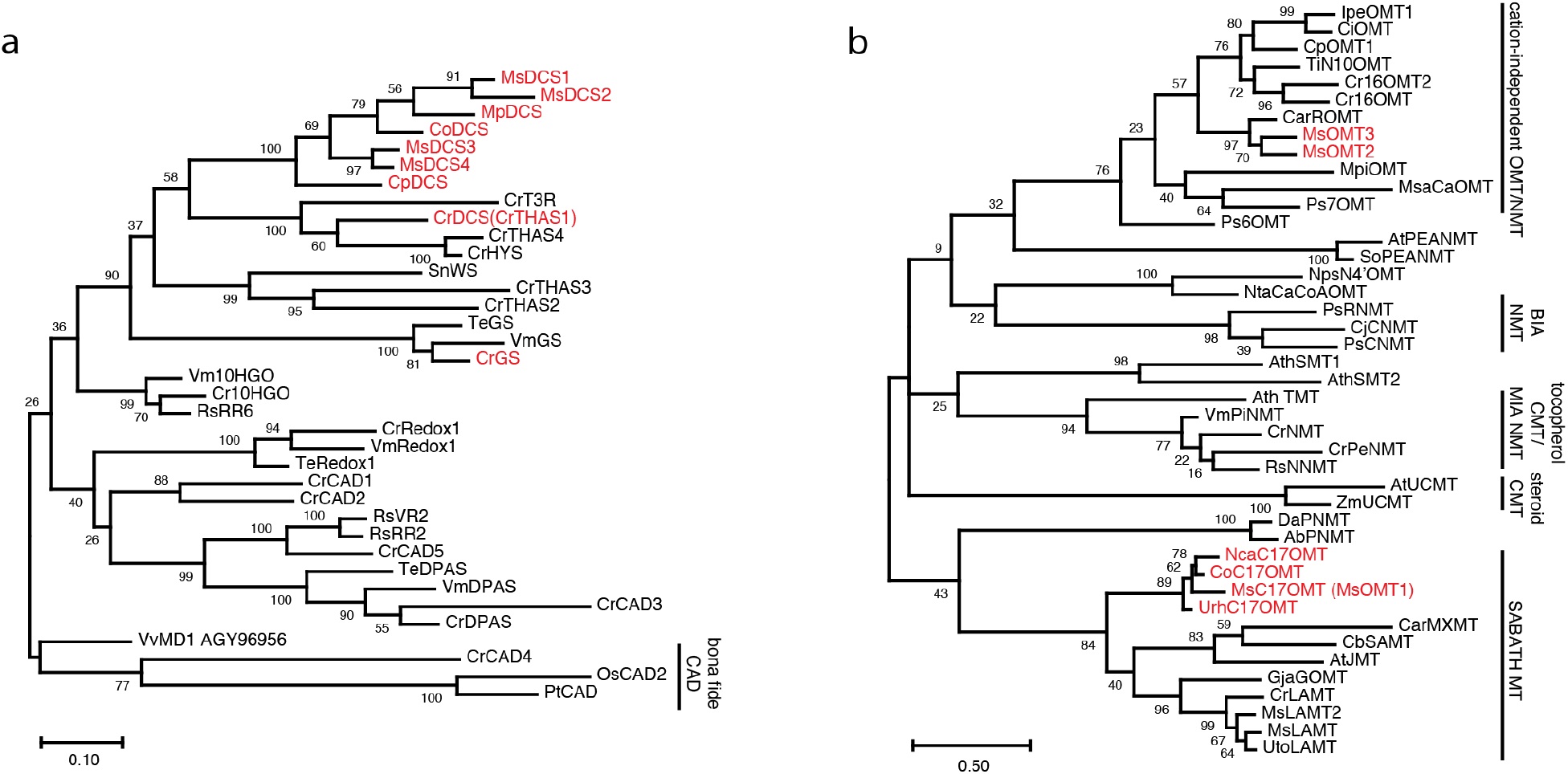
Phylogenetic analyses of short/medium chain dehydrogenases (a) and methyltransferases (b) in this study. The Genbank accession numbers for the proteins and plant species are included in Supplementary data 1. The enzymes labeled red are investigated in this study. The alignments were generated with MUSCLE algorism and the phylogenetic trees were generated with MEGA 10 (Maximum Likelihood method, Poisson correction model, 200 bootstrap replicates). 10HGO: 10-hydroxygeraniol oxidase; CAD: cinnamyl alcohol dehydrogenase; DCS: demethyldihydrocorynantheine/demethylcorynantheidine synthase; DPAS: dihydroprecondylocarpine acetate synthase; GS: geissoschizine synthase; HYS: heteroyohimbine synthase; THAS: tetrahydroalstonine synthase; VR2: vomilenine reductase 2; WS: Wieland-Gumlich aldehyde synthase; MT: methyltransferase.

We expressed and purified *N*-terminal His-tagged MsDCS1-4 in *E. coli*, and tested their activities in vitro. The strictosidine aglycones (*m/z* 351) were produced by the *C. roseus* strictosidine synthase (CrSTR) and strictosidine β-glucosidase (CrSGD) using the substrate tryptamine and secologanin (Fig. 1), which were coupled to MsDCS in vitro reactions. We also compared these reactions to those with the characterized enzymes CrGS, CrTHAS1, and CpDCS. As expected from previous studies, CrGS reduced strictosidine aglycones (*m/z* 351) to produce both 19*E*- and 19*Z*-geissoschizine (*m/z* 353, **3, 4**) with THA (*m/z* 353, **6**) by-product (Fig. 3). CpDCS produced a major product of demethyldihydrocorynantheine (*m/z* 355, 20*S*, **1**), which is intermediate to mitragynine 20*S*-epimer speciogynine (**17**) (Fig. 3). A number of isomeric MIAs (*m/z* 353 and *m/z* 355) were also produced as minor products in these reactions. We also noticed that CrTHAS1 produced not only THA but significant amounts of **1**, which has not been reported (Fig. 3). Similar to CpDCS, MsDCS2-4 produced **1** as a major product with other *m/z* 353 such as THA (**6**) and *m/z* 355 minor products (Fig. 3). In contrast, a major MsDCS1 product was a new *m/z* 355 MIA (Fig. 3) with an elution time different from that of **1**, which we tentatively identified as demethylcorynantheidine (20*R*, **2**). In the same reaction, MsDCS1 also produced 19*Z*-geissoschizine (*m/z* 353, **3, 4**), THA (**6**), yohimbine (*m/z* 355, by comparing to an authentic standard), and several other minor *m/z* 353 MIA products (Fig. 3).

**Figure 3.**
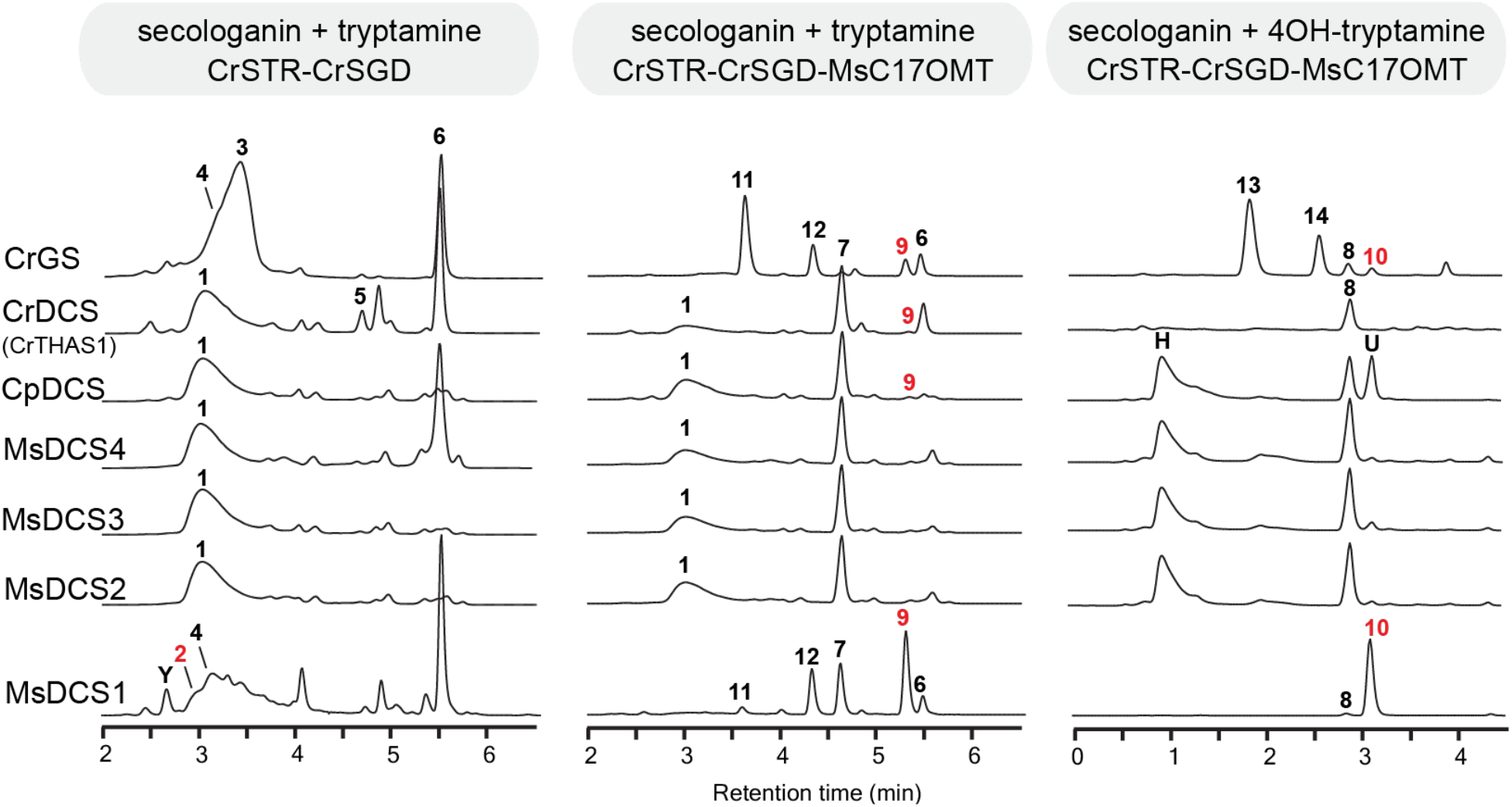
LC-MS/MS chromatograms of the coupled enzyme activities from CrSTR, CrSGD, CrGS, CrDCS (CrTHAS1), CpDCS, MsDCS1-4, and MsC17OMT (MsOMT1), with secologanin and tryptamine/4-hydroxytryptamine substrates. The peaks labeled with a red number indicate that they are intermediate to mitragynine. All MIA compounds were simultaneously recorded with *m/z* 353→144, 355→144, 367→144, 369→144, 383→160, and 385→160 ion transitions. The structures of peaks with a numeric number can be found in Figure 1. Peak Y: yohimbine; Peak H: 9-hydroxy-demethyldihydrocorynantheine; Peak U: unknown MIA with *m/z* 383.

With these findings, we continued to discover the methyltransferase (MT) responsible for 17-enol methylation. To our knowledge, no MTs discovered to date methylate a non-aromatic enol oxygen. The well-studied plant MTs firstly include the SABATH (**<u>S</u>**alicylic **<u>a</u>**cid carboxyl methyltransferase, **<u>B</u>**enzoic **<u>a</u>**cid carboxyl MT, and **<u>Th</u>**eobrobine synthase) MTs that methylate carboxylic acids such as salicylic acid and jasmonic acid ^13^. The loganic acid MT (LAMT) in *C. roseus* involved in MIA biosynthesis is also a member of this group ^14^. Other characterized MTs include the aromatic *O*-methyltransferases (OMT) that methylate the aromatic hydroxyls such as flavonoids and benzylisoquinoline alkaloids, steroid *C*-methyltransferases (CMT) that methylate many triterpenoids, and the tocopherol CMTs, some of which have evolved to perform MIA *N*-methylation ^15–18^ (Fig. 2b). Using the sequences of *C. roseus* CrLAMT and 16-hydroxytabersonine OMT(Cr16OMT, aromatic OMT) in MIA biosynthesis, we identified three candidate MTs from kratom transcriptome, which included a SABATH-like MT MsOMT1 and two homologous (70% amino acid identity) aromatic OMT type: MsOMT2 and 3. We further identified other MsOMT1 homologs from the transcriptomes of *Uncaria rhynchophylla* (cat’s claw), *Nauclea cadamba* (burflower tree), and *Cephalanthus occidentalis* (buttonbush), which all produce MIAs and reside in the Naucleeae tribe of the Rubiaceae family. No MsOMT1 homologs of more than 55% identity could be identified outside this tribe when searching in National Center for Biotechnology Information (NCBI) total non-redundant protein database, nor such homologs could be identified in another Rubiaceae species *Cinchona puberscens* in the tribe Cinchoneae or in the MIA producing species *C. roseus, Vinca minor*, and *Tabernaemontana elegans* in the Apocynaceae family. The phytogenic analysis (Fig. 2b) showed that MsOMT1 and its Naucleeae homologs form a distinct subgroup within the SABATH group, and they are clearly distinguished from other OMTs where MsOMT2 and 3 locate.

We expressed and purified *N*-terminal His-tagged MsOMT1-3 in *E. coli*, and tested their activities using purified 19*E*-, 19*Z*-geissoschizine (**3**,**4**), and demethyldihydrocorynantheine (20*S*, **1**). Clearly MsOMT1 was the expected C17-enol OMT, as it methylated all three substrates to respective geissoschizine methyl ethers (**11, 12**) and dihydrocorynantheine (**7**). The identity of 19*E*-geissoschizine methyl ether was also supported by comparing it to a commercial standard.

MsOMT1 showed typical Michaelis-Menten enzyme kinetics for all three substrates (Km 5.3-28.7 μM, Fig. 4). MsOMT1 did not show activity with other tested MIAs including reserpic acid, yohimbic acid, yohimbine, corynanthine, 3-hydroxy-2,3-dihydrotabersonine, 16-hydroxytabersonine, vincamine and ajmaline, or simple phenolic acids including gallic acid, syringic acid, salicylic acid, benzoic acid, caffeic acid, p-coumaric acid, ferulic acid and trans-cinnamic acid. The results suggested that MsOMT1 likely only accepts corynanthe MIA with a free 17-enol. We therefore renamed MsOMT1 as corynanthe 17-OMT (MsC17OMT). In comparison, MsOMT2 and 3 did not accept any tested MIA as substrates.

**Figure 4.**
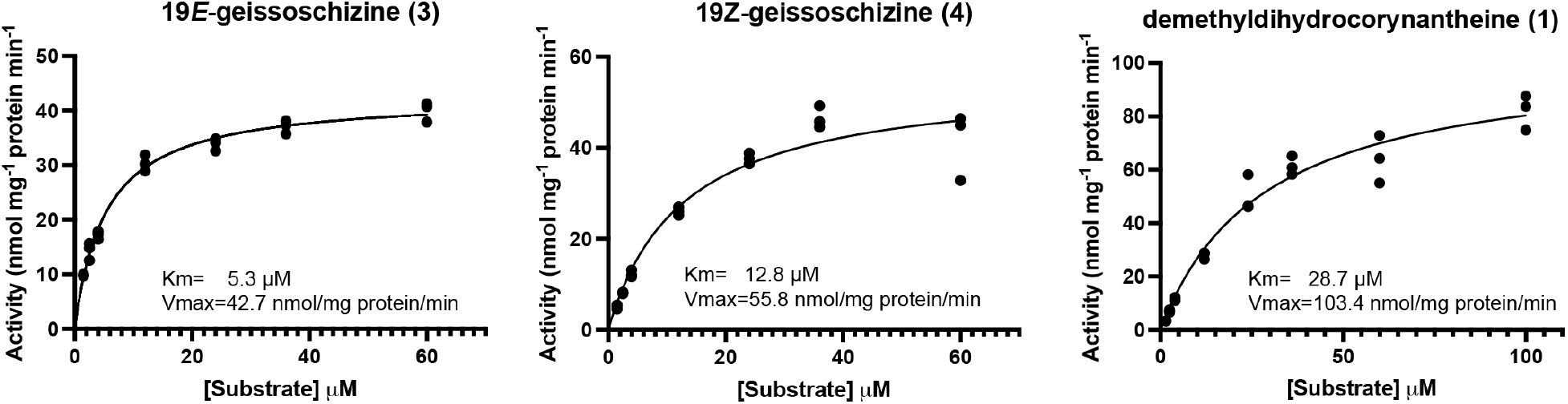
Michaelis-Menten enzyme saturation curves for MsC17OMT with three substrates: 19*E*-geissoschizine **(3**), 19*Z*-geissoschizine (**4**), and demethyldihydrocorynantheine (**2**). Data was generated with triplicates (_•_) for each substrate concentration.

With MsC17OMT, we revisited the activities of MsDCS1-4 in coupled reactions involving these enzymes. We confirmed that MsDCS2-4 produced a major product demethyldihydrocorynantheine (20*S*, **1**), which was methylated by MsC17OMT to dihydrocorynantheine (**7**) (Fig. 2). By 17-*O*-methylation, the reaction involving MsDCS1 clearly showed three major products: corynantheidine (20*R*, **9**) that is intermediate to mitragynine (**18**), dihydrocorynantheine (**7**), and 19*Z*-geissoschizine methyl ether (**12**) (Fig. 2). The most abundant product corynantheidine (**9**) was confirmed by comparing it to a commercial standard, which also confirmed our previous identification of demethylcorynantheidine (**2**) when MsC17OMT was not involved. With this finding, we further cloned two homologous reductases CoDCS and MpDCS from the plants buttonbush and *Mitragyna parvifolia*. When assayed in the same coupled reactions with MsC17OMT, both CoDCS and MpDCS led to the two products dihydrocorynantheine (**7**) and corynantheidine (**9**) (Supplementary Fig. 1). It is also worth noting that CrGS, CrTHAS1, and CpDCS also produced various amounts of corynantheidine (20*R*, **9**) in these MsC17OMT-coupled assays (Fig. 3), which could not be easily identified without 17-*O*-methylation due to the overlapping LC chromatograms of the non-methylated MIAs. These results indicated the plasticity of strictosidine aglycone reduction by these homologous reductases, which is key to the earlier diversification of MIA skeletons.

The remaining steps of mitragynine biosynthesis includes 9-hydroxylation and 9-*O*-methylation. We reasoned that the 9-hydroxyl group could be artificially introduced earlier in the biosynthesis by using 4-hydroxytryptamine substrate and the promiscuity of the biosynthetic enzymes. Despite with reduced activity, purified recombinant CrSTR, CrSGD, MsDCS1, and MsC17OMT converted secologanin and 4-hydroxytryptamine to demethylmitragynine (*m/z* 385, 20*R*, **10**) and demethylspeciogynine (*m/z* 385, 20*S*, **8**) epimers. We obtained similar results when we replaced MsDCS1 with MpDCS, while the assays containing MsDCS2-4, CrTHAS, CoDCS, and CpDCS only made demethylspeciogynine (*m/z* 385, 20*S*, **8**) as a major product (Fig. 3, Supplementary Fig. 1). Interestingly, CpDCS also produced a less reduced MIA (*m/z* 383, peak U in Fig. 3) that we have not identified. When we swapped MsDCS1 with GS, the reaction led to the formations of 9-hydroxy-geissoschizine 19-*E*/*Z* epimers (**13, 14**) that have not been reported in nature, as well as small amounts of both demethylmitragynine and demethylspeciogynine (Fig. 3). When we swapped the 4-hydroxytryptamine with 5, and 6-hydroxylated tryptamine in the same reactions, we detected the formations of 10, and 11-hydroxylated isomers of demethylmitragynine and demethylspeciogynine (Supplementary fig. 2), suggesting that both the reductases and MsC17OMT tolerate indole hydroxylations.

To confirm the identifications of demethylmitragynine (**10**) and demethylspeciogynine (**8**) from our in vitro reactions, we subjected these in vitro produced MIAs to the total leaf proteins isolated from kratom, buttonbush, *C. roseus*, and *Hamelia patens* (firebush), which were also supplemented with the methyl donor *S*-Adenosyl methionine (SAM). While the leaf proteins of kratom, buttonbush, or *C. roseus* did not catalyze the final 9-*O*-methylation, *H. patens* leaf proteins converted demethylmitragynine and demethylspeciogynine to the final products mitragynine (**18**) and speciogynine (**17**) that accumulate naturally in kratom leaves, when compared to authentic commercial standards (Fig. 5). The results again confirmed the identifications of other MIA intermediates, and indicated that the final methylation is SAM-dependent.

**Figure 5.**
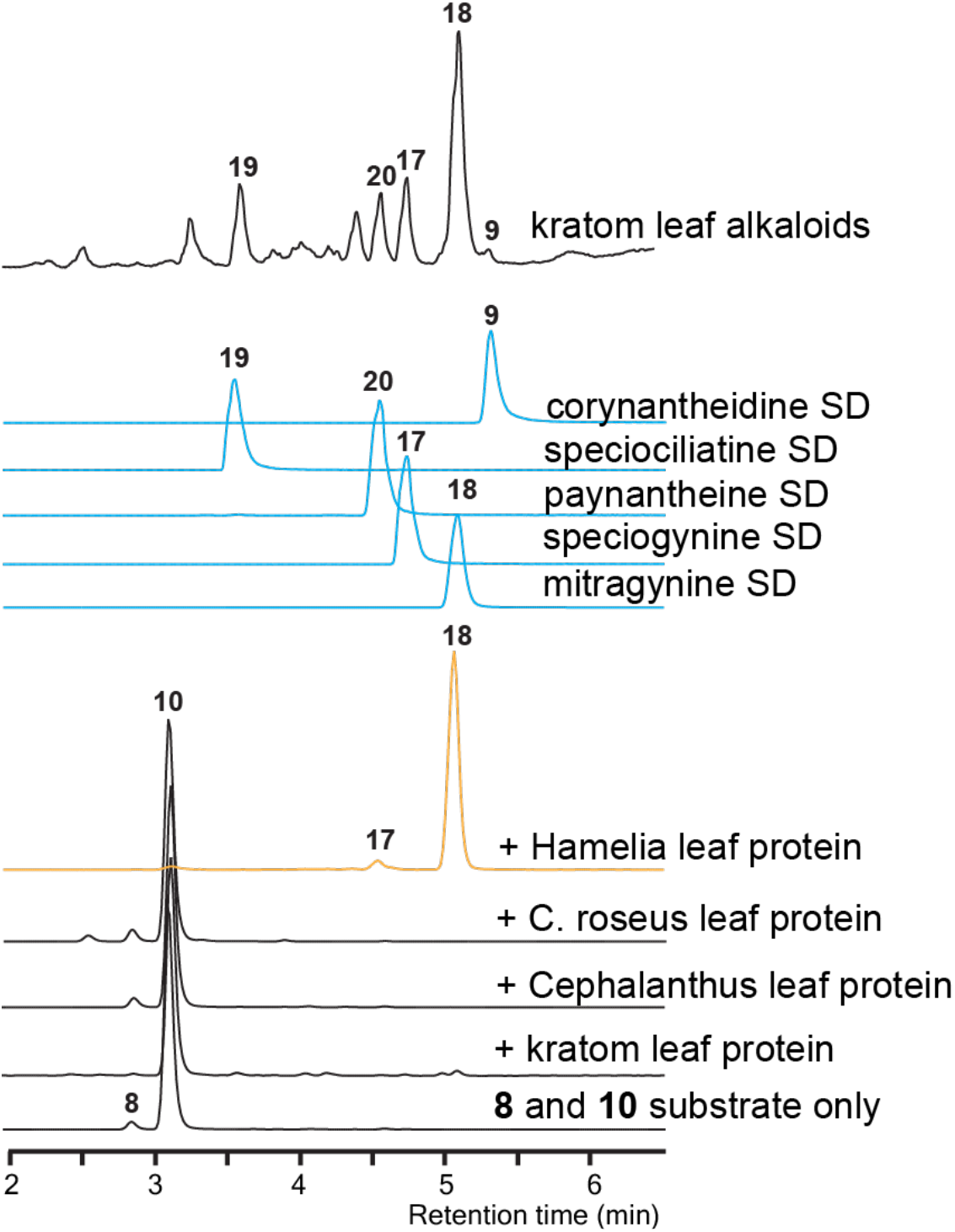
Demethylmitragynine (**10**) and demethylspeciogynine (**8**) were methylated to form mitragynine (**18**) and speciogynine (**17**) by a SAM-dependent methyltransferase from the total leaf proteins of *Hamelia patens*. All MIA compounds were simultaneously recorded with *m/z* 353→144, 355→144, 367→144, 369→144, 383→160, 385→160, 397→144, and 399→174 ion transitions by LC-MS/MS, except for kratom leaf alkaloids which were recorded with total ion chromatogram. Five alkaloid standards were used to identified in vitro enzyme products and the total kratom leaf alkaloids. The structures can be found in Figure 1.

## Discussion

Kratom alkaloids such as mitragynine, 7-hydroxymitragynine, and derivatives have been shown to be opioids that causes significantly less side effects and superior safety profiles ^1–5^. It is interesting that a pharmacophore based on corynanthe MIA skeleton can induce agonism with several opioid receptors, and such pharmacophore is amenable to further modifications that may lead to even safer opioids ^5^. The structure activity relationship studies shows that the 9-oxygen, 17-methoxyl, and the stereochemistry of C15 and C20 are all required for opioid receptor activations. For example, switching the C20 stereochemistry from R (mitragynine) to S (speciogynine) causes significantly loss of opioid activity ^2^, indicating the importance of this position in receptor binding. Therefore, it is critical to identify the biosynthesis responsible for these structural features. The biosynthesis of kratom opioids described here complements the recent invention of their total organic synthesis, and reveals an alternative method for acquiring these promising pharmaceuticals that may be developed commercially to mitigate the opioid overdose epidemics around the world.

In this study, we identified and characterized several homologous reductases that reduce the strictosidine aglycones to the corynantheidine (20R) and dihydrocorynantheine (20S) skeletons, as well as the critical C17-enol methyltransferase. Both enzymes are instrumental in forming the signature chemical structures that are required for kratom’s opioid activities. The identifications of the demethyldihydrocorynantheine/demethylcorynantheidine synthases (DCSs) were based on previous characterizations of these short/medium chain dehydrogenase types of reductases from *C. roseus* and *Cinchona puberscens* ^7–10^. The recombinant enzymes clearly showed great plasticity in their reduction product spectra since every enzyme was able to produce multiple products, which is owing to their peculiar structures and the well-known structural diversity of the strictosidine aglycones that exist in equilibrium (Fig. 1). Such product diversity was not easily discerned due to the low abundance of some products and their overlapping liquid chromatography behaviours. Remarkably, the C-17 enol methylation led to clear distinctions of these isomeric products, since the structures could no longer exist in enol-keto resonance. This allowed us to properly identify and characterize these strictosidine aglycone reductions, and identify the alternative products such as the demethylcorynantheidine (**2**) and demethyldihydrocorynantheine (**1**) products also produced by CrGS and CrTHAS1 (Fig. 2), which have not been described in previous studies. Based on the high production of demethyldihydrocorynantheine (**2**), we also suggest to rename CrTHAS1 as CrDCS.

Particularly, the SABATH type methyltransferase MsC17OMT was able to methylate the C17-enol on a corynanthe skeleton, which was demonstrated by using the geissoschizine 19-epimers, and demethylcorynantheidine 20-epimers (Fig. 3 and supplementary figs. 1 and 2). To our knowledge, this is the first example of enzymatic methylation of a non-aromatic enol in plant natural products. Other members of the SABATH methyltransferases included the founding members that methylate the carboxyl groups of plant hormones such as salicylic acid and jasmonic acid, as well as theobromine synthase that *N*-methylates 7-methylxanthine. The loganic acid methyltransferases (LAMTs) that perform the carboxyl methylation of MIA precursor loganic acid also belong to this group (Fig. 2b). The identification and characterization of MsC17OMT therefore demonstrated a new and notable member in the SABATH methyltransferase family. We could only find C17OMT homologs in the plants from the Naucleeae tribe of the Rubiaceae family, such as *Uncaria rhynchophylla* (cat’s claw), *Nauclea cadamba* (burflower tree), and *Cephalanthus occidentalis* (buttonbush). These plants naturally accumulate C17-methylated corynanthe MIAs and their simple derivatives such as the corresponding oxindoles, and are not known to rearrange the corynanthe skeletons to complex MIA skeletons, such as strychnos, aspidosperma, or iboga types that are commonly found in MIA producing plants in the Apocynaceae plants ^12,19,20^. Geissoschizine in *Catharanthus roseus* and *Rauwolfia serpentina*, both from the Apocynaceae family, is rearranged to strychnos skeleton (stemmadenine) and sarpagan skeleton (polyneuridine aldehyde) owing to the C17-enol reactivity and the cytochrome P450 monooxygenases that catalyze the intramolecular nucleophilic attack of the C17-enol to the iminium ^8,9,21^. It is likely that the C17OMT evolved in these Naucleeae species to mask the reactive C17-enol, which is otherwise prone to non-specific nucleophilic addition to other cellular components, as a way to reduce their cytotoxicity. However, such hypothesis requires further experimental confirmation.

While the 9-hydroxylase and 9-OMT that complete the mitragynine opioids biosynthesis still need to be characterized, we showed that the promiscuity of the pathway enzymes could accommodate the unnatural substrates: 4, 5, and 6-hydroxylated tryptamine and produce the corresponding 9, 10, and 11-hydroxylated corynanthe skeletons (Fig. 3 and Supplementary figs. 1and 2). We also showed that MsC17OMT could also accept these unnatural substrates and synthesize the corresponding C17-methyl ethers (Fig. 3 and Supplementary figs. 1 and 2). This enzyme promiscuity allowed us to synthesize demethylmitragynine (**10**) and demethylspeciogynine (**8**) by the activities of CrSTR, CrSGD, MsC17OMT and a number of DCS enzymes and GS from kratom, *M. parvifolia, C. occidentalis*, and *C. roseus*. Their structures of intermediates were confirmed multiple times with authentic standards along their biosynthesis towards the final products mitragynine (**18**) and speciogynine (**17**). We could not detect the 9-*O*-methylation activities using the leaf proteins from kratom, which might be caused by the protein extraction method that could not preserve the OMT activity. However, the 9-*O*-methylation of demethylmitragynine (**10**) and demethylspeciogynine (**8**) to mitragynine (**18**) and speciogynine (**17**) by *Hamelia patens* leaf total proteins and SAM methyl donor indicated that this step is a SAM-dependent methylation and a 9-OMT can also be identified in *Hamelia patens* that reside in the Hamelieae tribe of Rubiaceae family. The identifications of the 9-hydroxylase and 9-OMT will complete the mitragynine biosynthesis, which will lead to their complete biosynthesis without the needs of plant source or organic synthesis. Based on the enzyme promiscuity demonstrated in our study, it is likely the remaining enzymes may tolerate or could be engineered to accept other indole substitutions for enzymatic production of kratom opioid derivatives.

## Material and methods

### Chemicals standards

The standards for mitragynine, speciogynine, speciociliatine, paynantheine, corynantheidine, 4-hydroxytryptamine, and 5-hydroxytryptamine were purchased from Cayman Chemicals (Ann Arbor, MI, USA). The standard for 6-hydroxytryptamine was purchased from Toronto Research Chemicals (Toronto, ON, Canada), and the standard for 19*E*-geissoschizine methyl ether was purchased from AvaChem Scientific (San Antonio, TX, USA). The substrate 19E-geissoschizine and 19Z-geissoschizine was prepared and purified as described previously ^7^. The substrate demethyldihydrocorynantheine was prepared from in vitro reaction (10 mL) containing 20 mM Tris-HCl pH 7.5, 1 mM NADPH, 100 μg trptamine, 100 μg secologanine, and purified recombinant proteins: CrSTR (200 μg), CrSGD (20 μg), and MsDCS4 (200 μg). The reaction took place at 30 °C for 1 h, then it was extracted with 10 mL ethyl acetate. The evaporated extract was reconstituted in methanol and separated by thin layer chromatography using TLC-silica gel 60 f254 (Sigma-Aldrich) with solvent ethyl acetate : methanol (9:1, v:v), which afforded 20 μg demethyldihydrocorynantheine.

#### Plant materials, crude protein isolation, and RNA/cDNA synthesis

The plants *Mitragyna speciosa, Mitragyna parvifolia, Cephalanthus occidentalis, Cinchona puberscens, Catharanthus roseus*, and *Hamelia patens* were grown in a greenhouse at 28 °C with 16/8h photoperiod. Leaf tissues (3 g) and 0.2 g polyvinylpolypyrrolidone were ground in liquid nitrogen with mortar and pestle, which were extracted with ice-cold sample buffer (20 mM Tris-HCl pH 7.5, 100 mM NaCl, 10% (v/v) glycerol). The extracts were centrifuged at 15,000 g for 30 min, and desalted into the same sample buffer with a PD10 desalting column (ThermoFisher Scientific) according to the manufacture’s protocol. The total proteins were desalted one more time, and the final samples were stored at -80°C. Leaf tissues (100 mg) were collected for RNA extraction using standard TRIzol RNA isolation reagent according to the manufacture’s protocol (ThermoFisher Scientific). The resulting RNA was used to generate cDNA using the LunaScript®RT SuperMix Kit according to the manufacture’s protocol (New England Biolabs).

#### Cloning

From respective leaf cDNA, MsDCS1, MsDCS2, and MpDCS were amplified using the same primer set (1/2). MsDCS3 and MsDCS4 were amplified using the same primer set (3/4). CpDCS was amplified using the primer set (5/6). CoDCS was amplified using primer set (7/8). MsC17OMT (MsOMT1), MsOMT2, and MsOMT3 were amplified using primer set (9-14) respectively. The primers are listed in Supplementary table 2. MsOMsDCS1-4, MpDCS, MsC17OMT, MsOMT2 were cloned in pET30b+ vector within BamHI/SalI site, and mobilized to E. coli BL21DE3 for expression. MsOMT3 was cloned in pET30b+ vector within SalI/NotI site, and mobilized to E. coli BL21DE3 for expression. CpDCS and CoDCS were gateway-cloned into pDEST17 vector by Gateway™ BP and LR clonase™ II Enzyme mix according to manufacturer’s protocol (ThermoFisher Scientific), and mobilized to E. coli BL21A1 for expression.

### Recombinant protein expression and purifications

An overnight culture (2 ml) of E. coli BL21DE3 strains containing MsDCS1-4, MpDCS, MsC17OMT, MsOMT2, and MsOMT3 in pET30b+ vectors were used to inoculate 200 ml LB media, which were cultured at 200 rpm and 37°C until OD600 reached 0.6-0.7. The cultures were induced with 0.1 mM IPTG at 15°C, 200 rpm overnight. For CpDCS and CoDCS, an overnight culture (2 ml) of E. coli BL21A1 was used to inoculate 200 ml LB media, which were cultured at 200 rpm and 37°C until OD600 reached 0.3. The cultures were induced with 0.1% (w/v) galactose at 15°C, 200 rpm overnight. The induced cultures were sonicated in ice-cold sample buffer (20 mM Tris-HCl pH 7.5, 100 mM NaCl, 10% (v/v) glycerol) and purified using standard Ni-NTA affinity chromatography. After eluting with 250 mM imidazole in sample buffer, the purified recombinant proteins were desalted using a PD-10 desalting column (GE Health Sciences) according to manufacturer’s protocol into the same sample buffer and stored at

-80°C.

### In vitro assays and kinetics

A standard in vitro reaction (50 μl) included 20 mM Tris-HCl pH7.5, 1 mM NADPH, and some or all these components: 60 μM SAM, 2 μg CrSTR, 0.2 μg CrSGD, 2 μg MsDCS1-4, CoDCS, MpDCS, CrGS, or CrDCS(CrTHAS1), and 2 μg MsC17OMT. The substrates included 1 ug secologanine and 1 μg tryptamine or its hydroxylated derivatives. The reaction was incubated at 30°C for 1 h and terminated by adding 150 μL methanol. The kinetics triplicated assays (50 μl) included 20 mM Tris-HCl pH7.5, 100 μM SAM, 1 μg MsC17OMT, and substrate 19*E*-, or 19*Z*-geissoschizine, or demethyldihydrocorynantheine at 1.5, 2.5, 4, 12, 24, 36, 60 and 100 μM concentrations. The kinetics assays were performed at 30°C for 2 min before they were terminated by adding 150 μL methanol to the reactions. The products were quantified using a standard curve, to generate the enzyme velocity. The kinetics parameters and saturation curves were approximated using the software Prism 9.5.0 (GraphPad Software, LLC.).

### LC-MS/MS

LC-MS/MS was performed on an Agilent Ultivo Triple Quadrupole LC-MS equipped with an Avantor® ACE® UltraCore™ SuperC18™ column (2.5 μm, 50×3mm), which included the solvent systems: solvent A, methanol: acetonitrile: ammonium acetate 1 M: water at 29:71:2:398; solvent B, methanol: acetonitrile: ammonium acetate 1 M: water at 130:320:0.25:49.7. The following linear gradient (8 min, 0.6 ml/min) were used: 0 min 80% A, 20% B; 0.5 min, 80% A, 20%B; 5.5 min 1% A, 99% B; 5.8 min 1% A, 99% B; 6.5 min 80% A, 20% B; 8 min 80% A, 20% B. The photodiode array detector records from 200 to 500 nm. The MS/MS was operated with gas temperature at 300°C, gas flow of 10 L/min, capillary voltage 4 kV, fragmentor 135 V, collision energy 30V with positive polarity. The Qualitative Analysis 10.0 software by Agilent was used for all LC analyses.

