## Supplementary information for "Biosynthesis of Kratom Opioids"

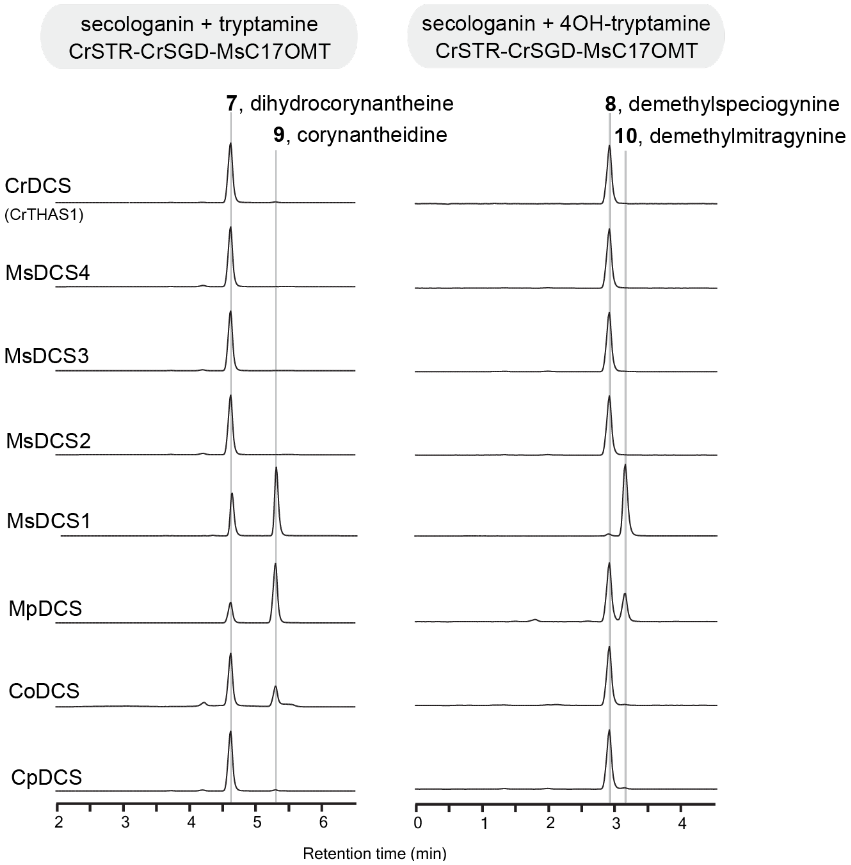


**Supplementary figure 1.** LC-MS/MS chromatograms of the coupled enzyme activities from CrSTR, CrSGD, CrDCS (CrTHAS1), CpDCS, MsDCS1-4, MpDCS, CoDCS and MsC17OMT (MsOMT1), with secologanin and tryptamine/4-hydroxytryptamine substrates. Selected MRM ion transitions of *m/z* 369→144 and 385→160 were used for left panel and right panel, respectively. Ms: *Mitragyna speciosa*; Mp: *Mitragyna parvifolia*; Co: *Cephalanthus occidentalis*; Cp: *Cinchona puberscens*; Cr: *Catharanthus roseus*. The structures can be found in Figure 1.

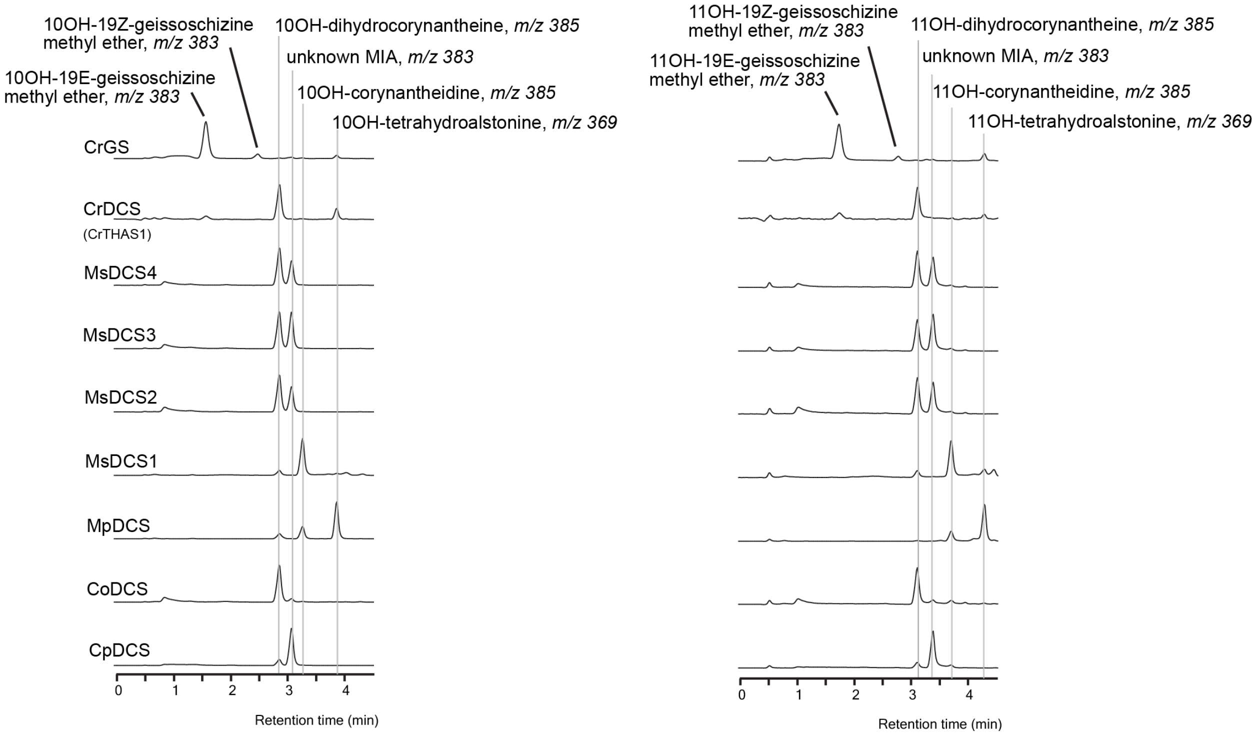


**Supplementary figure 2.** LC-MS/MS chromatograms of the coupled enzyme activities from CrSTR, CrSGD, CrGS, CrDCS (CrTHAS1), CpDCS, MsDCS1-4, MpDCS, CoDCS and MsC17OMT (MsOMT1), with secologanin and 5-hydroxytryptamine/6-hydroxytryptamine substrates. All MIA compounds were simultaneously recorded with *m/z* 353→144, 355→144, 367→144, 369→144, 383→160, and 385→160 ion transitions. Ms: *Mitragyna speciosa*; Mp: *Mitragyna parvifolia*; Co: *Cephalanthus occidentalis*; Cp: *Cinchona puberscens*; Cr: *Catharanthus roseus*. The structures can be found in Figure 1.

Supplementary table 2. Primers used in this study.

| # | primer name | primer sequence 5'-3' |
|---|---|---|
| 1 | MsDCS1-BamHI-F | TTAGGATCCTGAAATGGCAGGAAAATGTCCTC |
| 2 | MsDCS1-SalI-R | ATAGTCGACTTAAGGTGATTTCAGTGTATTCCCG |
| 3 | MsDCS3-BamHI-F | TTAGGATCCAATGGCAGGAAAATCTCCCGAA |
| 4 | MsDCS3-SalI-R | ATAGTCGACAAGCAACTTTAAGAAGTTGCA |
| 5 | Attb-CpDCS-F | GGGGACAAGTTTGTACAAAAAAGCAGGCTTCATGGCCGGAAAATCTCAAGAAG |
| 6 | Attb-CpDCS-R | GGGGACCACTTTGTACAAGAAAGCTGGGTCAAGCAGATTTTAAGGTATTGCC |
| 7 | Attb-CoDCS-F | GGGGACAAGTTTGTACAAAAAAGCAGGCTTCATGGCAGGAAAATCTCCCCAAG |
| 8 | Attb-CoDCS-R | GGGGACCACTTTGTACAAGAAAGCTGGGTTAAGCTGATTTCAGTGTATTCCCG |
| 9 | MsOMT1-BamHI-F | TTAGGATCCAATGCAACCACAGAGAGGGAG |
| 10 | MsOMT1-SalI-R | ATAGTCGACTCAATCCTCTGCATTGCGTTTA |
| 11 | MsOMT2-BamHI-F | TTAGGATCCAATGGATTTGGCTAGAAATGGT |
| 12 | MsOMT2-SalI-R | ATAGTCGACTCAATAATAGATCTCAATGAGGG |
| 13 | MsOMT3-SalI-F | ATAGTCGACAGCATGGATTTGGCTAGAAAAAG |
| 14 | MsOMT3-NotI-R | AGCGGCCGCGGTATGAAACAATTCAAGTACATCA |
